# The control costs of human brain dynamics

**DOI:** 10.1101/2024.01.24.577068

**Authors:** Eric G Ceballos, Andrea I Luppi, Gabriel Castrillon, Manish Saggar, Bratislav Misic, Valentin Riedl

**Affiliations:** Montréal Neurological Institute, McGill University, Montréal, QC, Canada; Department of Psychiatry and Behavioral Sciences, Stanford University, Stanford, CA, USA; Department of Neuroradiology, Klinikum rechts der Isar, TUM School of Medicine and Health, Technical University of Munich, Munich, Germany; Department of Neuroradiology, Uniklinikum Erlangen, Friedrich-Alexander-University Erlangen-Nuremberg, Erlangen, Germany; Research Group in Medical Imaging, SURA Ayudas Diagnósticas, Medellín, Colombia

## Abstract

The human brain is a complex system with high metabolic demands and extensive connectivity that requires control to balance energy consumption and functional efficiency over time. How this control is manifested on a whole-brain scale is largely unexplored, particularly what the associated costs are. Using network control theory, here we introduce a novel concept, time-averaged control energy (TCE), to quantify the cost of controlling human brain dynamics at rest, as measured from functional and diffusion MRI. Importantly, TCE spatially correlates with oxygen metabolism measures from positron emission tomography, providing insight into the bioenergetic footing of resting state control. Examining the temporal dimension of control costs, we find that brain state transitions along a hierarchical axis from sensory to association areas are more efficient in terms of control costs and more frequent within hierarchical groups than between. This inverse correlation between temporal control costs and state visits suggests a mechanism for maintaining functional diversity while minimizing energy expenditure. By unpacking the temporal dimension of control costs, we contribute to the neuroscientific understanding of how the brain governs its functionality while managing energy expenses.

## INTRODUCTION

The intricate nature of the human brain, characterized by its extensive connectivity [5, 10, 81] and correspondingly high metabolic demands [58, 65], has led to the assumption that its inherent organization represents a delicate balance between energy consumption and functional efficiency [70]. This notion underscores the role of the nervous system as a regulatory entity [27], orchestrating its functions to fulfill cognitive demands while simultaneously managing energy-related constraints. While prior research has shed light on how the brain manages its energy expenditures at a cellular level [28, 43, 58], the broader exploration of how these mechanisms translate to the regulation of functional costs at a whole-brain scale remains relatively uncharted.

Drawing inspiration from engineering principles, network control theory (NCT) offers a novel perspective on this problem by conceptualizing the brain as a networked control system in order to explain its dynamics [33]. In its most basic form, NCT considers brain dynamics as a composite outcome of a region’s connectivity profile and the necessary control inputs to guide neural activity toward a desired state [33, 39, 60] (Fig. 1a). The former aspect delves into the constant anatomical interactions between various brain regions, while the latter presents an adaptable measure to optimally transition between states within the confines of energy constraints (Fig. 1b). These constraints are quantified as control energy, which measures the costs entailed in controlling the brain across diverse states (Fig. 1c).

**Figure 1.**
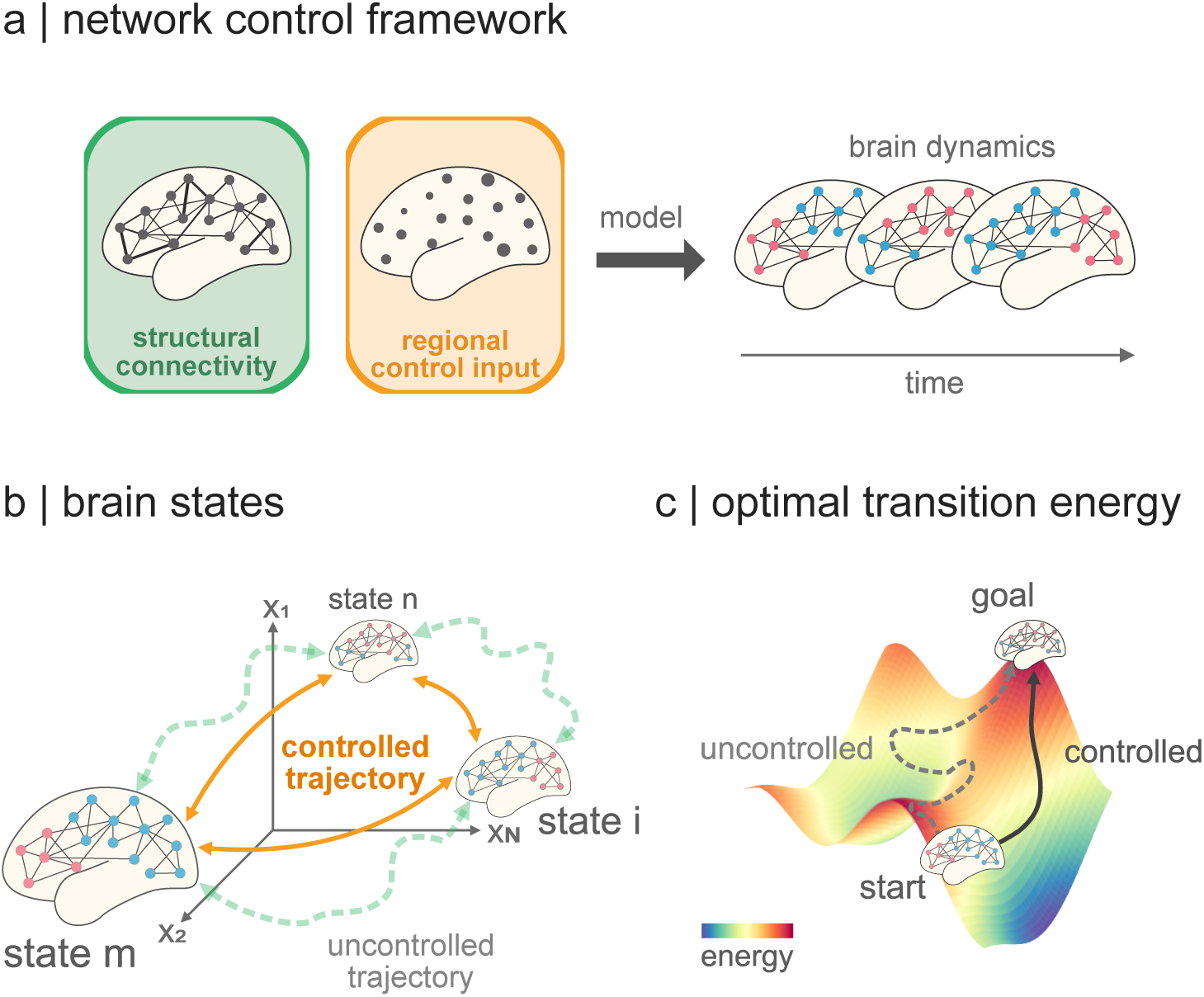
Network control theory. (a) The brain as a network of connected regions fluctuates through different modes of activity across time. We can model these transitions throughout time using network control theory. Network control theory is a framework that uses information about the anatomical connections in the brain, which are fixed, and regional control inputs, which are optimised, as bases to model the temporal evolution of brain activity (highlighted by the changing node color patterns on the right). In other words, the changes in brain activity of a particular region are modelled as depending on its connectivity profile and an “activation” that spreads out along such profile, but also spreads in from other regions. Such activation is referred to control input and is fitted to represent an optimal bridge between the brain’s current and next state [60]. (b) Before employing a control model, we define what a brain state is, i.e., how maps of brain activity are represented. In previous literature, states have been defined as functional intrinsic networks [15, 17, 36], cytoarchitectonical hierarchy levels [61], or cognitive maps [48], to name a few. These states are represented as single points in an *N* -dimensional state space, with *N* being the number of brain regions that we control. We seek to find the shortest possible trajectory between two points in this space and generate activations such that the brain is steered, or “controlled” along a previously optimised trajectory. (c) We define the current and next state of brain activity to be related by the structural connectome of an individual (see *Methods*). The connectome imposes a resistance, or flow, into the state space such that it is easier to make activity changes between states in one direction than another. Think of walking on a hill where the walk uphill will be more strenuous than the downhill walk. In analogy to this, it costs the brain more energy, called “control energy”, to control its activity between particular states than others. This asymmetry has been the subject of study in previous literature [48, 61]

Control energy has been shown to be a powerful tool in various domains of neuroscience, providing an explanation of how the enhancement of executive functions is supported by structural changes across development [17], explaining the effects of psychedelics on brain dynamics [71], or predicting cortical responses to electrical stimulation [79]. Nevertheless, studies employing NCT traditionally focus on the control energy between single pairs of states without accounting for how such costs organize in time. Furthermore, control energy remains a purely statistical notion [78], dissociated from the actual energetic currency employed by the brain (e.g., glucose and oxygen metabolism), with only recent efforts attempting to establish a connection between control costs and metabolism by showing that relative hemispheric differences in glucose uptake are mirrored by differences in control energy for temporal lobe epilepsy patients [36].

Despite these advances, the costs associated with controlling neurotypical resting state activity as well as their energetic signature remain elusive. In light of these considerations, we propose a novel framework to examine the expenses incurred in regulating resting state dynamics, i.e., activity over time. Building upon the findings of previous reports [36], we further investigate whether theoretically derived control costs manifest in tangible biological metrics of energy expenditure by comparing them to normative measures of energy metabolism. Moreover, we extend our methodology to investigate organizational principles in brain dynamics through control costs. Ultimately, our efforts seek to establish a foundation for comprehending how the brain governs its functionality and the associated costs entailed in this intricate control.

## RESULTS

Here, we present our methodology to study the control costs of human brain dynamics, i.e., activity changes over time, using network control theory. We begin by describing our analysis pipeline and the modelling choices made to simulate the regional energy expenditure based on anatomical and functional properties from diffusion and functional magnetic resonance imaging (MRI) data from *n* = 327 participants in the Human Connectome Project Young Adult dataset [87]. Our work introduces a new measure of control energy over time, time-averaged control energy (TCE), which we relate to actual measurements of energy metabolism based on positron emission tomography (PET). Finally, we conclude our results by investigating the transition costs between functional hierarchical levels in the brain and find a dynamical rule whereby brain switch frequency within and between levels of the cortical hierarchy is inversely proportional to its control cost.

### Simulating the control costs of resting state dynamics

We take a multi-step approach to simulate the control costs of human brain dynamics. In brief, our procedure consists of identifying the dominant networks at each time point (Fig. 2a). This information is used to estimate the average activity across time-points where each network dominates (Fig. 2b) in order to create state maps for each intrinsic network (Fig. 2c). We then use the sequence of network dominance to track the control energy required to transition through the respective states (Fig. 2d). Finally, we average across all transitions, yielding an estimate of the average costs to control brain activity over time (Fig. 2e).

**Figure 2.**
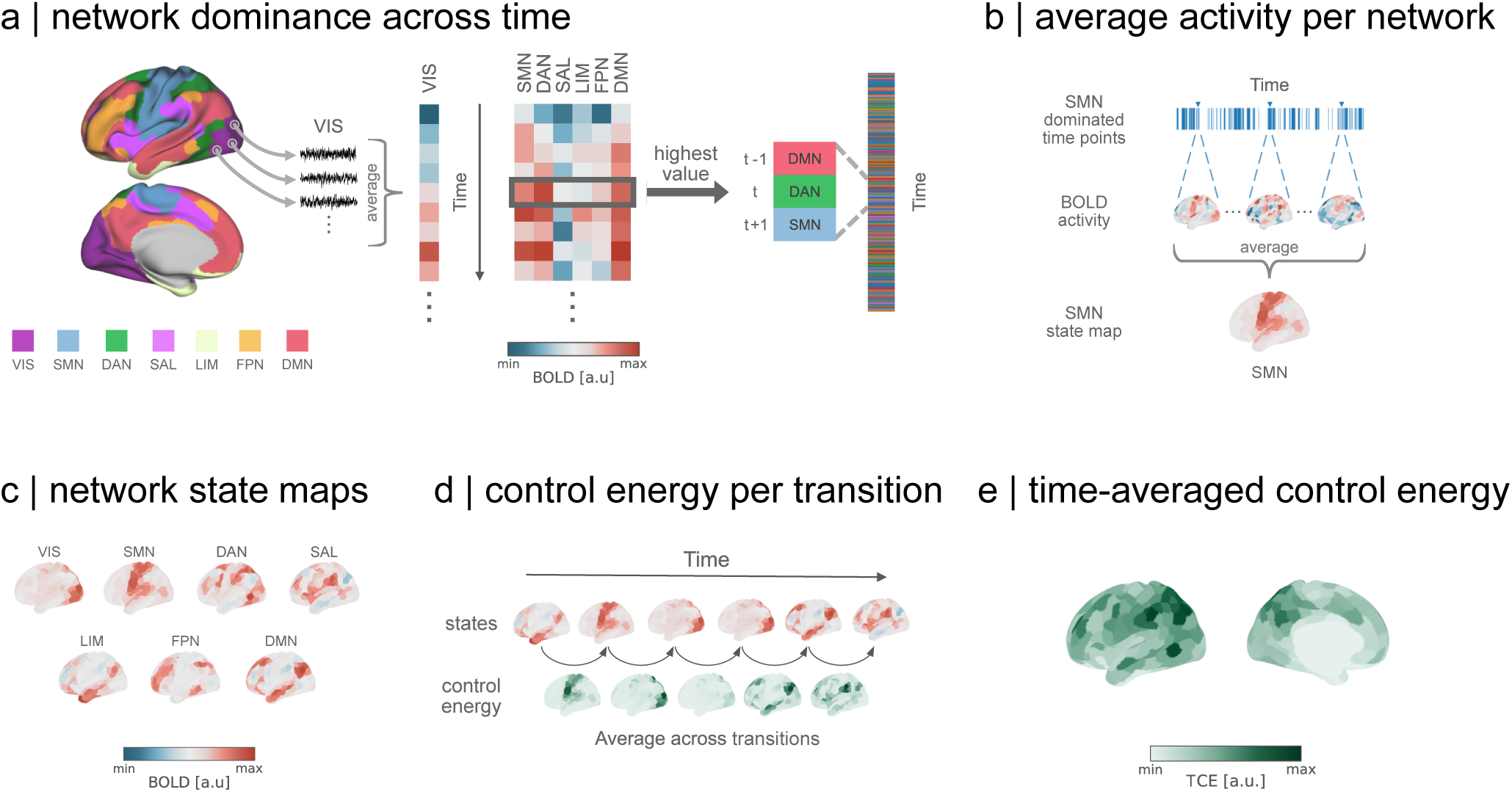
Estimating the costs of brain dynamics. (a) We begin by averaging the activity of all regions belonging to one of seven canonical intrinsic networks of Yeo-Krienen [95] in the Schaefer-parcellation [69] and compare the amplitude in each time point. For every time point, the network with the highest amplitude is assigned as the dominating network in that moment, yielding a sequence of network dominance across the time (right). (b) Using the time markers of dominance for each network, BOLD activity maps are averaged to generate a representative state map of activity. (c) We generate seven representative network state maps, reflecting the average magnitude and sign of cortical activity during each network activation. (d) Following the sequence of network dominance, we simulate the optimal control strategy in which the brain transitions between pairs of network states. Thus, each transition yields a map of momentary control costs to move between states. We finally average all control maps across transitions. (e) Our framework results in a map of time-averaged control energy for each individual, where the value of each region represents the amount of control energy integration a region saw throughout the recording on average. VIS: visual network; SMN: somatomotor network; DAN: dorsal attention network; SAL: salience ventral network; LIM: limbic network; FPN: frontoparietal network; DMN: default mode network; BOLD: blood-oxygen-level-dependent; TCE: time-averaged control energy.

Our analysis begins by delineating intrinsic networks as brain maps. When seeking to model task-related brain states, it is possible to rely on statistical maps of parameter estimates [8] or meta-analytic maps [48]. However, neither of these approaches is feasible for modelling brain activity at rest. Previous studies have taken two approaches to model resting states: On the one hand, states have been defined according to a priori specified parcellations, where areas that fall into a given intrinsic network are represented with ones and all other areas with zeros [17, 36]. On the other, resting states have been derived from clustering techniques and then categorized based on their similarity to binary intrinsic network maps like in the previous approach [15, 71].

While both approaches offer a respective advantage, they also come with their drawbacks. The former approach profits from the interpretability of previously established brain networks with defined neurocognitive functions. However, such binary, discontinuous state representations are equally defined for all individuals, therefore losing important unique information about the subject. The latter approach circumvents this by using datadriven clustering techniques to derive states from individual data. However, given the possibility of multiple solutions in clustering [23, 37, 41], there is no guarantee that any representation of intrinsic networks are present, thereby complicating the interpretability and replicability of findings across studies. We thus propose a new approach that preserves the intelligibility of intrinsic networks as states, is non-binary, and tailored to individual data.

Our approach relies on an a priori definition of intrinsic networks, such as the canonical intrinsic networks proposed by Yeo-Krienen [95], which are widely-used in the neuroimaging literature. We begin by partitioning the voxel-wise BOLD imaging data into parcel time series representing the time-activity of a region from an intrinsic network. The time series are then z-scored across time and later averaged with other time series from the same network (Fig. 2a). This yields a network-wise average activity for each time point in the recording. Intrinsic networks are then compared at each time point based on their average activity and the network with the highest activity is designated the dominating network of that time point. To control for noise, we discard time points where no network activity surpasses a fixed threshold set at half a standard deviation in average activity. We redo our analysis with a lower threshold and also leave out the network modelling altogether to validate our approach (see *Sensitivity and robustness analysis*)

With the identified time points of dominance for each intrinsic network in hand we proceed to compute the actual state representations of networks. The dominance labels serve as temporal markers to define which time points belong together. We average the activity within such markers to yield a centroid-like representation that resembles an intrinsic network, with a continuous magnitude and sign that is captured by the activity of each individual (Fig. 2b). These state coefficient maps represent each intrinsic network in the a priori defined parcellation, with additional information about regions outside the network that co-fluctuate with it (Fig. 2c). Essentially, we are defining network representations for each individual by tailoring the network weights to their brain activity.

To simulate the control energy between each network coefficient map, we leverage structural information about the anatomical connections of each individual, derived from diffusion MRI. As referenced earlier, network control methods require state maps to represent initial and target positions from where we simulate the control costs to move between them. Such costs are subject to anatomical constrains imposed by the individual’s connectome. In other words, we quantify the effort required in each region to change its current activity to a desired state, with other connected regions also contributing to its change [33, 39, 60]. To maintain biological plausibility, we set our simulated path to follow an optimal trajectory that balances the energy required to enable a transition with the proximity to the desired target state [33]. More simply, we simulated transitions that focus on energy minimization while imposing constraints to prevent them from deviating into implausible, non-physiological pathways that may otherwise provide a low-energy pathway.

The results of our modelling of pairwise state transitions manifest in an asymmetric matrix representing the energy cost required to transition the entire brain from one network state to another (Suppl. Fig. S2). The inherent asymmetry in these costs arises from the directional flow imposed by the structural connectivity within the state space. In another context, the states we have modeled could be thought of lying on peaks and valleys, where access is significantly easier in one direction than the other — a phenomenon analogous to the contrast between ascending and descending a hill, with the former requiring more energy.

Having captured the control cost of the transition between all pairs of resting states, we now reconcile this information with the temporal sequence of the dominant networks. This means that we use the network sequence to simulate the energy required for the transitions between states as they occurred during the recording (Fig. 2d). We aggregate the transition energy over all pairs of states in the recording sequence and then normalize the cumulative energy by dividing it by the number of transitions, i.e., we average the overall energy across transitions. This normalization ensures that our measurement remains comparable across different recording durations. Ultimately, our approach culminates in the derivation of a metric we refer to as time-averaged control energy, or TCE (Fig. 2e).

TCE is an estimate of the temporal control costs associated with idiosyncratic intrinsic network activations across time (Fig. 3a). It is a spatially heterogeneous measure but we find that its distribution in left and right hemispheres is statistically indistinguishable [Mann-Whitney U = 76390, P = 0.27](Fig. 3b). Moreover, previous evidence indicates that control energy is modulated by age [17, 80]. We find that stratifying by age (using bins of 22-25, 26-30, 31-35, and 36+, with 247, 527, 418 and 14 individuals, respectively) does not lead to any statistically significant group differences [Levene W = 108, P = 0.955; ANOVA F (3, 1596) = 0.133, P = 0.94] (Fig. 3c). Taken together, we show that the control costs of human brain dynamics are regionspecific, consistent between hemispheres, and conserved across ages studied here.

**Figure 3.**
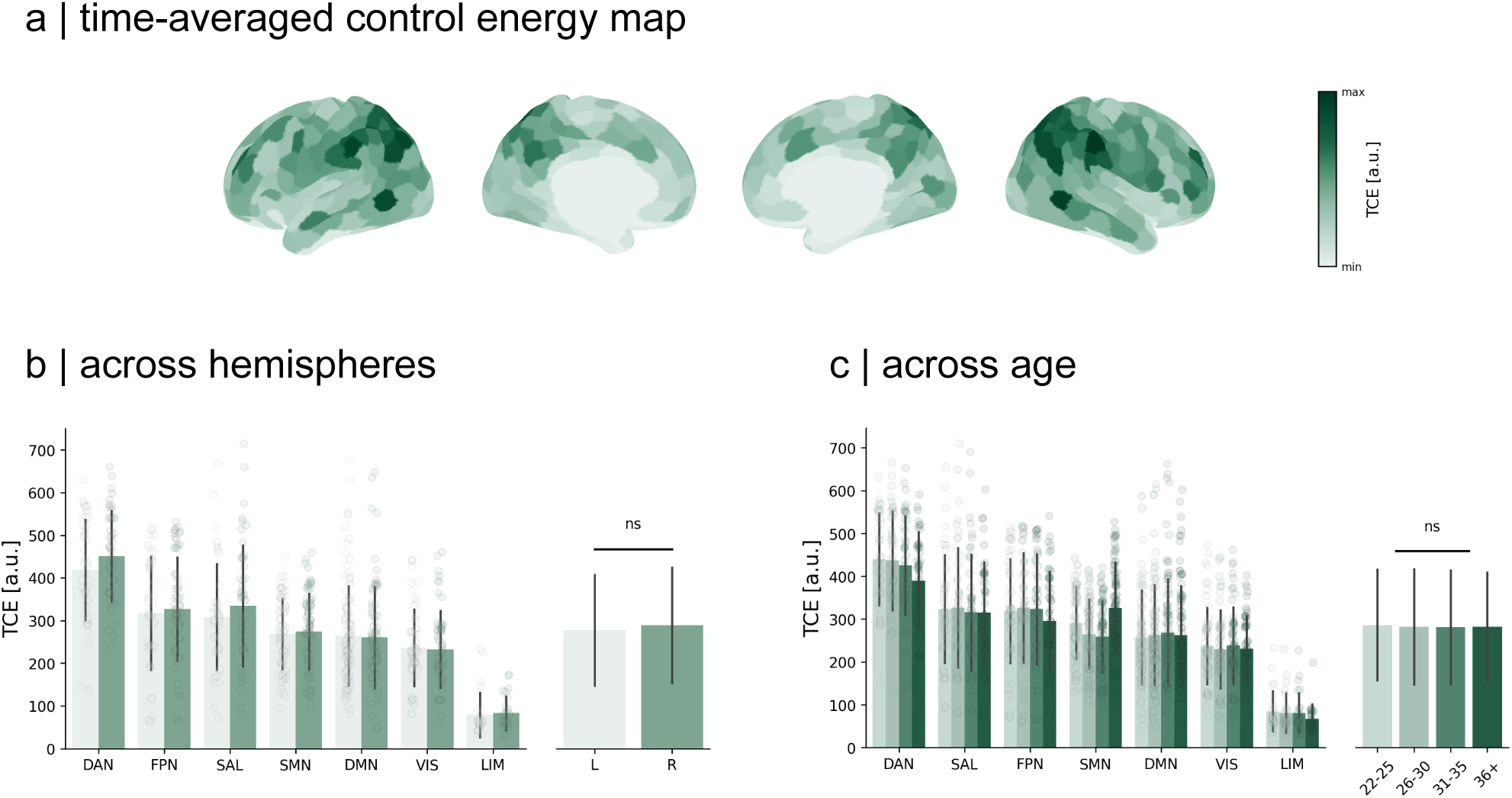
Time-Averaged Control Energy. (a) Averaging all TCE maps across individuals, we observe that the costs to control resting state dynamics are regionally heterogeneous. (b) Regions in the dorsal attention network require the most control with limbic regions costing the least. In general, temporal control costs display no preference for a hemisphere, with comparable values evident in both the left and right hemispheres [Mann-Whitney U = 76390, P = 0.27]. (c) When stratifying our analysis by age, we see no statistical difference in temporal control costs across ages groups [ANOVA F(3, 1596) = 0.133, P = 0.94]. Each dot represents the subject-averaged value for a region-of-interest. Bars around the mean indicate standard deviation of the data.

### Grounding control costs in metabolism

Previously, network control theory metrics were shown to correlate with physiological markers of energy metabolism [36], thereby establishing an initial bridge between theoretical and tangible indicators of energy. As such, we contextualize our results with neurobiological maps to look for energetic correlates of temporal control costs. To accomplish this, we scrutinize the relationship between temporal control costs and two pivotal substrates of metabolic energy: oxygen and glucose metabolism [13, 65] (Fig. 4).

**Figure 4.**
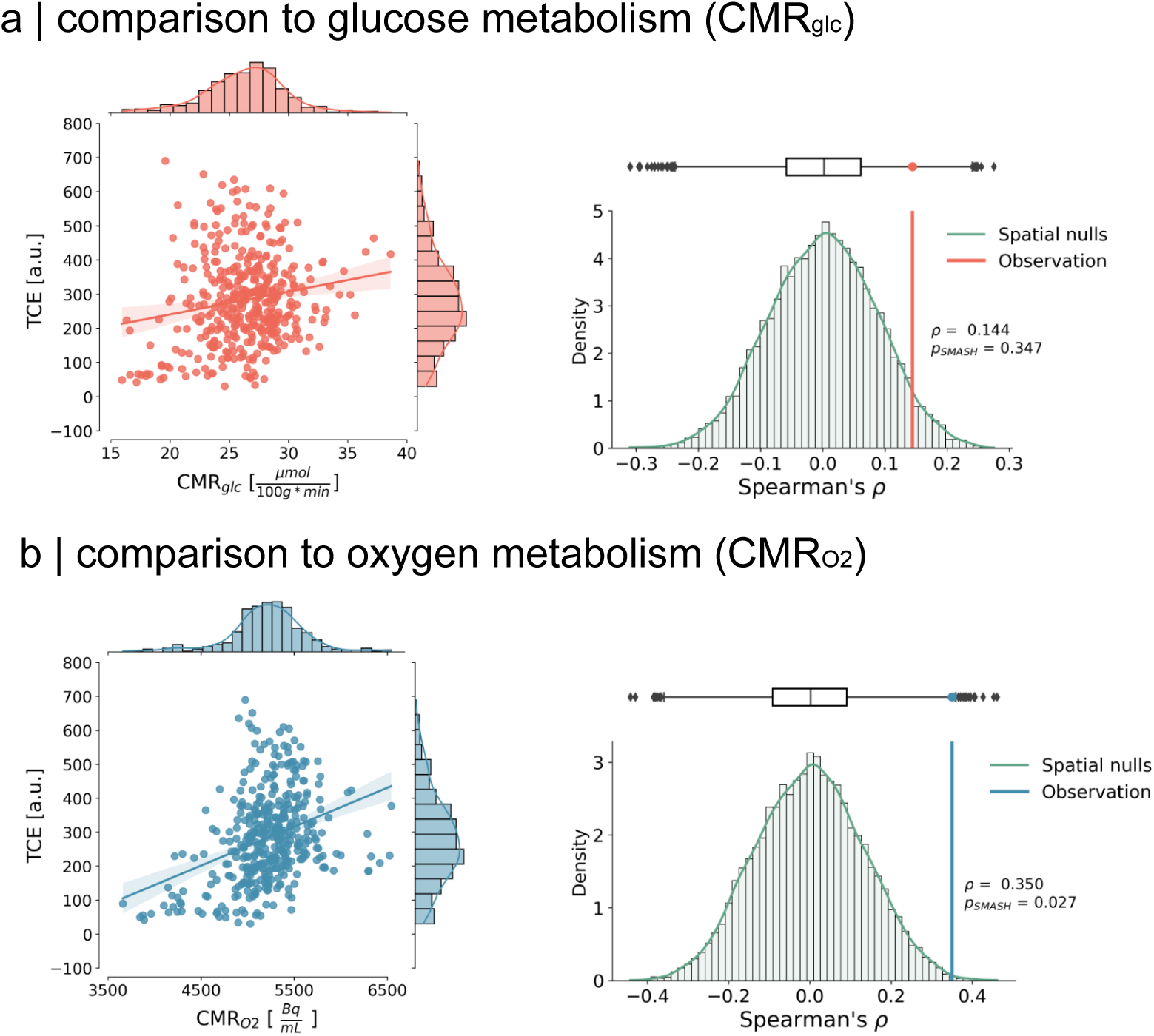
Relationship to metabolism. (a) Left: We compared how time-averaged control energy spatially co-varies with glucose metabolism from an internally acquired positron emission tomography (PET) dataset [14]. Glucose uptake has been previously shown to scale with momentary control energy [36]. We, however, do not find a similar significant relationship to control energy when considering its temporal dimension [Spearman *ρ* = 0.144, *PSMASH* = 0.347]. Shades around regression line represent 95% confidence interval with 1000 bootstrap samples. Right: Distribution of spatial nulls. Correlation results accounted for spatial autocorrelation in the data by using variogram-matched nulls [12, 51, 88]. (b) Left: Contrary to glucose metabolism, we found that TCE is spatially similar to a representative cortical distribution of oxygen uptake from PET imaging [86] [Spearman *ρ* = 0.35, *PSMASH* = 0.027]. Shades around regression line represent 95% confidence interval with 1000 bootstrap samples. Right: Distribution of spatial nulls. Correlation results accounted for spatial autocorrelation in the data by using variogram-matched nulls [12, 51, 88].

Our initial investigation involved comparing TCE to the regional variations in cerebral glucose metabolism (*CMR_glc_*) across the cortex. To achieve this, we employed a group-average template derived from an internally acquired dataset [14] comprising *n* = 20 subjects who underwent PET scans, utilizing ^18^F-Fluorodeoxyglucose (FDG) as a tracer to monitor their glucose uptake (see *Methods*).

Given the inherent spatial smoothness of our PET map, we employ rigorous measures to validate our results by contrasting them against 10,000 null maps that replicate the variogram of the data [12] (see *Methods)*. These null maps randomize the values in our PET map while accounting for the spatial autocorrelation inherent in the recordings, enabling us to distinguish true effects from spatially induced artifacts using a modified P-value (P*_SMASH_*) [12, 51]. Our results showed a weak correlation between TCE and *CMR_glc_*, which, however, did not exhibit statistical significance when the spatial smoothness of the data set was taken into account [Spearman *ρ* = 0.144, *P* = 0.003, *P_SMASH_* = 0.347].

Subsequently, we investigate the relationship between TCE and cerebral oxygen uptake within the cortex. Here, we employed a normative map of cerebral oxygen metabolism (*CMR_O_*_2_) derived from [86]. This map was computed from *n* = 33 participants who underwent PET scans to estimate their cerebral oxygen consumption, utilizing ^15^O-labeled oxygen (see *Methods*). We note that the map is not quantified into cerebral metabolic units; however, we retain the name for oxygen uptake as *CMR_O_*_2_, in accordance with the original authors’ nomenclature for their activity concentration.

Given the same technical limitations as with the *CMR_glc_* map, we generate 10,000 null maps that preserve the spatial autocorrelation of the original data using the same variogram-matching algorithm [12]. Our results show that TCE significantly correlates with *CMR_O_*_2_, and that this relationship is significant even after accounting for the spatial autocorrelation in the data [Spearman *ρ* = 0.35, *P* < 0.001, *P_SMASH_* = 0.027]. Altogether, we find that the regional cortical pattern of temporal control costs resembles the spatial pattern of oxygen uptake, establishing further evidence about the link between biological and control energy.

### Temporal organisation of control costs

We now turn to a new perspective on brain states by reframing our analysis from the perspective of functional hierarchical processing as defined by Mesulam [54]. Previous work has shown that control energy differs across cortical hierarchies [61]. In this context, we examine the control costs of switching within and between hierarchical levels across time, ultimately looking at their dynamical organisation in time.

We begin by mapping the sequence of dominant networks to their respective levels within the functional hierarchy of the brain (Fig. 5a). Visual and sensorimotor networks are hereafter referred to as “unimodal”, a distinction that arises from their specialized processing that assumes specific inputs[49, 54]. In turn, we refer to all other networks as “heteromodal” (associative) as they integrate multiple functional streams [54]. Bi-modal mapping ensures consistency in our subsequent validation with a different parcellation that defines intrinsic networks differently (Suppl. Fig. S5). Moreover, we show that adopting this new perspective provides a fresh view of the dynamics of the brain, uncovering a relationship between TCE and state switching behavior. We follow the same framework to compute TCE, but now separately average transitions within unimodal networks, i.e., from unimodal to unimodal, and within heteromodal networks, as well as transitions between unimodal and heteromodal levels.

**Figure 5.**
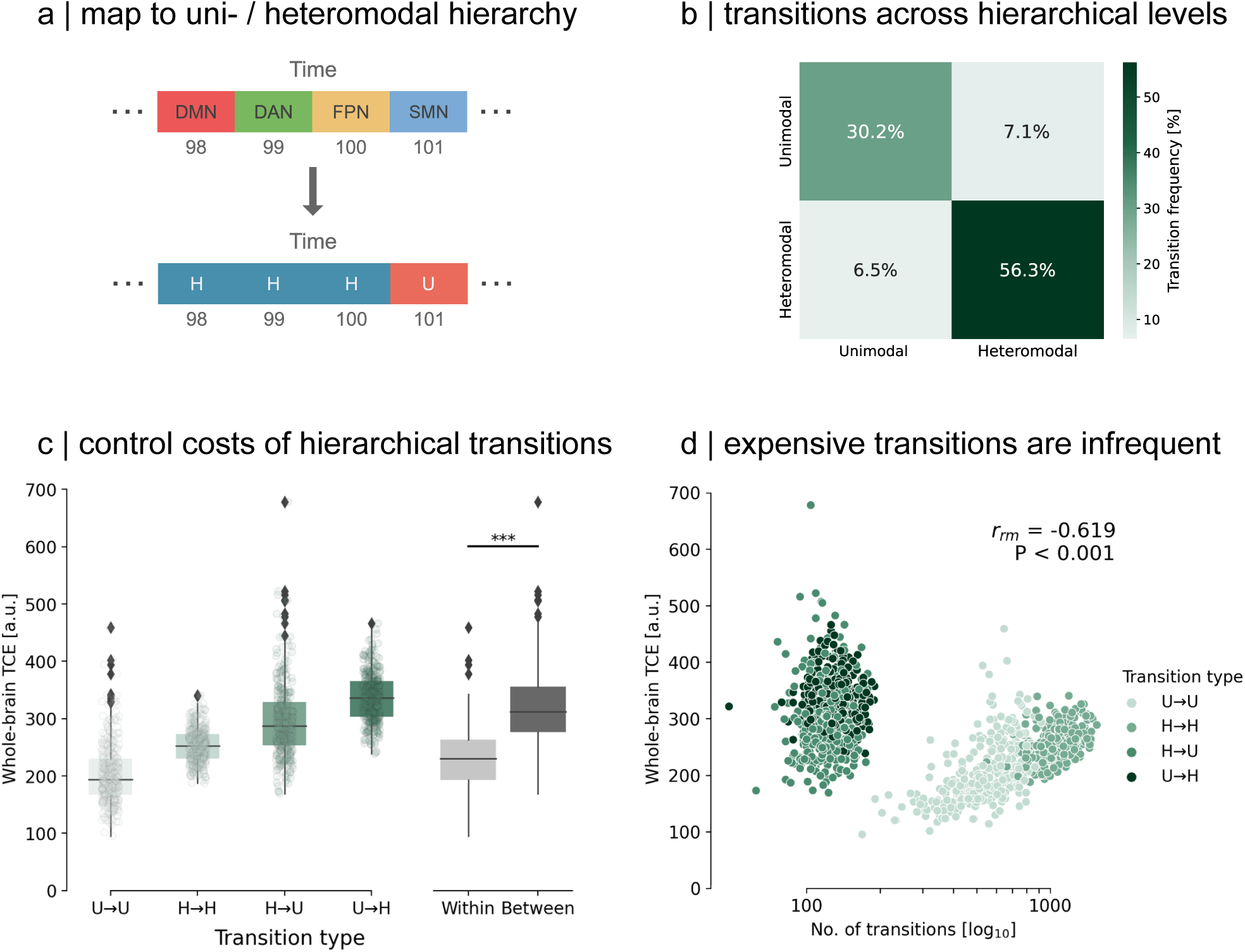
Temporal organisation of control costs. (a) We map our sequence of dominating networks to their respective hierarchical level. This means sensory networks are jointly labelled as unimodal networks (U), with the remaining networks defined as heteromodal, association networks (H). (b) Transitions within hierarchical levels represent the majority of transitions (86.5%) at rest. (c) Averaging control costs across the whole brain, we observed that transitions within a hierarchical level required significantly less TCE than between. Asterisks indicate high significance [Mann-Whitney U = 376072, P < 0.001]. Each dot per transition type represents a subject. (d) We relate whole-brain control costs to transition frequency using repeated measures correlation [4] to assess common intra-individual relationships and find that the number of transitions per transition type is negatively correlated to the cost needed to perform such switch [*rrm* =-0.619, P < 0.001].

After mapping the sequence of dominating networks, we observe that, on average for individuals, transitions occur predominantly within a hierarchy rather than between hierarchies (Fig. 5b). Analogously, the control costs to transition between hierarchies are more demanding than those within a hierarchical level [Mann-Whitney U = 376072, P < 0.001](Fig. 5c). Using repeated measures correlation [4, see *Methods*] to account for repeated measures of control costs for each participant as well as their individual traits, we examine the commonalities in the relationship between participants’ control costs and transition frequency across hierarchies. We find a robust negative correlation throughout the cohort [*r_rm_* =-0.619, P < 0.001], indicating that more frequent transitions within a hierarchy are associated with lower average control energy costs, in stark contrast to infrequent, costly transitions that bridge different hierarchical domains. We show that this relationship is not an artefact of the network partition in our parcellation, but is also evident when using alternative partitions (see *Sensitivity and robustness analysis*). Altogether, our findings suggests a dynamic temporal principle within the brain that strives to minimize its control costs by keeping expensive transitions sparse.

### Sensitivity and robustness analysis

To ensure the robustness of our results, we replicate our analyses on an independent dataset [14]. This additional dataset has lower spatial and temporal resolution and is subjected to different preprocessing protocols (see *Methods*). Indeed, even in the face of these differences, our results remain unchanged (Suppl. Fig. S6).

We then repeat our analysis using an alternative cortical parcellation [30]. This approach seeks to validate the reproducibility of our results with a different definition of intrinsic networks. We find that this approach leads to consistent results (Suppl. Fig. S5). Additionally, we investigate the impact of a coarser partition by dividing the cortex into 200 regions instead of the original 400 regions. We find that this modification does not affect our results (Suppl. Fig. S4).

Further, we test the impact of our state modelling step by leaving it out and instead simulate the control costs to transition between time points of raw activity, i.e., our state maps are represented as the BOLD activity at each time point. We form a group-average template like in our main results and compare these results to the temporal signal-to-noise ratio (tSNR) of each region. tSNR is a metric to quantify the quality of a BOLD signal, calculated as the mean of the signal prior to demeaning and divided by the signal standard deviation [92]. We compute the tSNR for each participant run in the HCP dataset, re-scale it from 0 to 1 using min-max normalization, and finally average it across runs.

We find that, when leaving out the state modelling step, regions with high TCE are also regions that exhibit a low temporal signal-to-noise ratio (tSNR) [Spearman *ρ* = 0.36, *P_spin_* = 0.042]. In contrast, using our approach we observe that the group-level results are unlikely related to the group-level regional tSNR [Spearman *ρ* = 0.234, *P_SMASH_*= 0.232]. Following our main approach, we discard time points with average activity below 0.5 but observe that, using a lower threshold of 0 instead, our group-level results again show no significant relationship to regional signal quality [Spearman *ρ* = 0.274, *P_SMASH_* = 0.099]. Moreover, we observed on an individual level that a larger extent of participants’ TCE maps were significantly related to their tSNR when leaving out our modelling approach (Fig. S1). Taken together, this suggests that modelling states using our approach mitigates the influence of noise in our analysis.

We conclude by examining whether temporal control costs are related to the occurrence of resting states. This would mean that the patterning of individual TCE maps are a reflection of the most frequently occurring states. To test this, we create a representative map of the occurring intrinsic networks by counting how frequent, i.e., how many time points, each state occurs. Dividing this number by the total number of time points yields an estimate of how much each network was present in the recording, e.g., the visual network dominated 20% of the time. We multiply this ratio with its corresponding state coefficient map, thereby creating time-weighted maps which we can sum together to create a map that represents the frequency of each network and its spatial distribution. Interestingly, we find no significant association between this map and our TCE map (Spearman *ρ* = 0.27, *P_SMASH_* = 0.092; Suppl. Fig. S3). Altogether, we conclude that our findings result from a convergence of characteristic sequences of resting states and individual anatomical constraints, which together form the basis to simulate the control costs of resting state dynamics.

## DISCUSSION

In the present study, we introduce a metric for assessing the control costs inherent in human brain dynamics, referred to as time-averaged control energy (TCE). TCE serves as a composite measure quantifying the average control exerted by an individual to sustain dynamic neural activity over time. Notably, our findings reveal a statistically significant spatial resemblance between the distribution of TCE and cortical oxygen metabolism, a major energetic substrate crucial for compensating neural activity [35]. Our results further show that transitions within unior heteromodal networks entail distinct temporal control costs as opposed to transitions between modalities, thereby offering insight into the temporal organisation of brain dynamics. Importantly, we provide open-access code to facilitate the replication and extension of our results in subsequent studies.

### Temporal control is a confluence of anatomical constraints and state sequences

Network control theory is a powerful tool to investigate the emergence of dynamics from the structural scaffold of the brain [60]. Its application in neuroscience has yielded valuable insights, e.g., into individual differences in psychiatric conditions [48, 62] or developmental changes in youth [17, 80]. Notably, controllability is reported to increase asymptotically and plateau with adolescence [80], potentially explaining the absence of age effects observed in our results.

We find that the distribution of temporal control costs is spatially distinct, with certain brain regions requiring higher TCE than others (Fig. 3). The extent of engagement in this control dynamic may be influenced by the manner in which control inputs propagate, reminiscent of the role of connectivity profiles in facilitating diverse spreading dynamics across brain regions [56]. This spreading process may parallel the flow of control throughout the connectome [78].

Control costs are further influenced by the specific states the brain traverses. Our results corroborate previous findings [48, 61] indicating that control energy is dependent on both initial and goal states (Fig. 5; Suppl. Fig. S2). This underscores the significance of our state- modelling approach that determines the sequence of network dominance based on intrinsic network activations, thereby establishing the order of state transitions.

Our methodology demonstrates robustness across different partitions of intrinsic networks (Suppl. Fig. S5, S4), as well as separate datasets (Suppl. Fig. S6). Nonetheless, our state-modelling approach trades off robustness for temporal granularity, thereby losing the ability to capture temporal shifts in brain activity. Indeed, brain dynamics are hypothesized to be temporally organised such that high-amplitude modes of activity are counterbalanced by transient ’off’ states [67, 90, 99]. We prompt future investigations to experiment with alternative state-modelling methods, like sliding-window approaches, for improved temporal precision. Appropriate tests should, however, be developed in tandem to distill real fluctuations from spurious ones [42, 97].

Finally, we note that our model of network control is linear, therefore assuming that linear dynamics govern brain transition [39]. While nonlinear models of neural dynamics have shown promise in simulating brain activity [9], the use of first-order linear approximations has proven sufficient to model activity at a macroscale [57]. Furthermore, recent studies have showcased how such linear control models are expandable to account for biological heterogeneities in receptor availability [71, 72] or cortical abnormalities in patient populations [48]. Consequently, we see potential for future efforts to expand on our work in order to investigate how temporal control costs are modulated by incorporating additional biological insights or clinical distinctions.

### Control costs reflect metabolic features of the brain

Our results show that temporal control costs colocalize with oxygen metabolism, but not glucose metabolism (Fig. 4). A previous study from He et al. [36] showed that control energy and glucose uptake are similarly put out of balance between hemispheres in temporal lobe epilepsy (TLE) patients. The authors, however, only show this relationship in the limbic system and, moreover, their methodology does not consider the costs of temporal switching, but seeks to link relative hemispheric asymmetries in momentary control costs and glucose uptake to explain structural changes in TLE patients. In contrast, our study relates control and metabolism on an absolute whole-brain basis and considers the temporal dimension of control costs.

Moreover, while both glucose and oxygen constitute fundamental components for meeting the energetic demands of neural activity [13, 35, 44, 53, 86, 91], they exhibit distinct spatial patterns. The predominant energy-producing pathways in the brain encompass glycolysis and oxidative metabolism, where glycolysis relies on glucose and oxidative metabolism relies on glucose-derived pyruvate and oxygen [13, 21]. The disparate spatial distribution of glucose and oxygen metabolism is attributed to their dissimilar upscaling capacities for energy generation [20, 22, 96]. Specifically, certain regions are posited to favor glycolysis beyond basal levels of oxygen and glucose consumption at an equable ratio in order to afford greater flexibility in energy demand necessary for supporting human cognition [47, 86]. In light of these considerations, our association of temporal control costs with the spatial distribution of oxygen, rather than glucose metabolism, supports the interpretation that control of human brain dynamics is grounded in baseline metabolic activity constituent of oxidative energy generation.

We note, however, that our conclusion is based on group-level comparisons of control costs and metabolic maps. Further research is warranted to investigate the nuanced relationship between control costs and energy metabolism at an individual level. Advances in MR imaging now enable the simultaneous acquisition of BOLD and oxygen metabolism data from the same subjects [24], providing an avenue for in-depth exploration of this relationship. Finally, we underline that our results are not confounded by the fact that we use BOLD- imaging data, as the BOLD signal reflects blood flow, which represents oxygen availability rather than necessarily indicating oxygen consumption [26].

### State switching is grounded by control costs

We observed that the costs required to transition within and between hierarchical levels, i.e., unimodal or heteromodal, were significantly different (Fig. 5c,d). Similarly, this phenomenon was mirrored in the frequency with which the brain either transitions out or persists in a hierarchical level (Fig. 5b,d). This means that costly transitions between hierarchical levels were rare, with more efficient transitions within hierarchical levels happening more frequently.

In order to enable information flow, the human brain is posited to switch between modes of integration and segregation over time, where segregation refers to regions clustering together to form functionally distinct modules and integration represents a form of global intercommunication between modules [18, 25, 70]. Building upon this, we propose that the transitions between functional hierarchical levels may reflect an integration-segregation dynamic. In this sense, switches between hierarchical levels might represent modes of communication among functional hierarchies, counterbalanced by switches within hierarchical levels as modes of segregated activity. As previous studies have shown the disruption of integration-segregation dynamics by pathology or altered states of consciousness [45, 46], future works could investigate whether such conditions equally disrupt state switching dynamics.

Furthermore, we hypothesize that the inverse correlation between control costs and state visits represents a mechanism where the brain seeks to maintain functional diversity, while minimizing its energy expenditure. This is similar to the hypothesis put forward by Zalesky, et al. [98], where the authors posit that the brain transitions through costly, intermittent states in order to facilitate global integration while keeping energy demand minimal. As such, we encourage future efforts to further elucidate our understanding of the intricacies governing state switching dynamics and control costs.

In conclusion, our work presents a new perspective of control energy as a measure of the costs to control human brain dynamics. We are able to show that temporal control costs are spatially related to oxygen metabolism as an energy substrate; and uncover an organisational principle that relates temporal dynamics to their costs in the brain. As such, we envision that our work can help advance research on the costs of regulating brain activity.

## METHODS

All preprocessed data is available at https://osf.io/nw9zt. The code and additional data used to perform the analyses are available at https://github.com/NeuroenergeticsLab/control_costs.

### Data

#### Human Connectome Project (HCP)

The main data used for this study consisted of resting state time series from functional MRI (fMRI) and structural connectomes from diffusion MRI (dMRI) taken from the Human Connectome Project S900 Young Adult release [87]. Scans from 327 unrelated participants (mean age 28.6 ± 3.73 years, 55% females) were used to ensure that familial factors do not confound our analysis [55]. Informed consent was obtained for all subjects (the protocol was approved by the Washington University Institutional Review Board as part of the HCP). The participants were scanned in the HCP’s custom Siemens 3T “Connectome Skyra” scanner, and the acquisition protocol included four 15-minute resting state fMRI sessions and a high angular resolution diffusion imaging (HARDI) sequence. The resting state fMRI data was acquired using a gradient-echo EPI sequence (TR = 720 ms; TE = 33.1 ms; FOV = 208 × 180 mm^2^; voxel size = 2 mm^3^; number of slices = 72; and number of volumes = 1200). The dMRI data was acquired with a spin-echo EPI sequence (TR = 5520 ms; TE = 89.5 ms; FOV = 210 × 180 mm^2^; voxel size = 1.25 mm^3^; b-value = three different shells i.e., 1000, 2000, and 3000 s/mm^2^; number of diffusion directions = 270; and number of b0 images = 18). Additional information regarding the acquisition protocol can be found under [87].

To process the functional data, each run of each subject’s resting state fMRI recording was pre-processed in terms of gradient distortion correction, motion correction, and spatial normalization according to [76] and [29]. Artefacts were then removed using ICA-FIX [68]. Inter-subject registration of cerebral cortex was carried out using areal feature-based alignment and the Multimodal Surface Matching algorithm [66]. The clean time series were then parcellated into 400 and 200 cortical region time series according to the Schaefer functional atlas[69].

To process the diffusion data, structural connectomes were reconstructed from the dMRI data using the MR- trix3 package [84]. Grey matter was parcellated into 400 and 200 cortical regions according to the Schaefer functional atlas [69] and fiber orientation distributions were generated using a multi-shell multi-tissue constrained spherical deconvolution algorithm [19, 38]. The initial tractogram was generated with 40 million streamlines, with a maximum tract length of 250 and a fractional anisotropy cutoff of 0.06. Spherical-deconvolution informed filtering of tractograms (SIFT2) was used to reconstruct whole brain streamlines weighted by cross-section multipliers [74]. More information regarding the individual network reconstructions is available in [59].

#### TUM Dataset

The replication data consisted of simultaneously measured FDG-PET and fMRI, with subsequent dMRI recordings, all while the participants kept their eyes open. Scans from 20 participants (mean age 34.2 ± 5.99 years, 50% females) were used. Informed consent was obtained for all subjects (the protocol was approved by the Technical University of Munich Review Board). The participants were scanned an integrated PET/MR (3T) Siemens Biograph mMR scanner (Siemens, Erlangen, Germany) and used a 12- channel phase-array head coil for the MRI acquisition. The PET data were collected in list-mode format with an average intravenous bolus injection of 184 MBq (s.d. = 12 MBq) of [18F]FDG. In parallel to the PET measurement, automatic arterial blood samples were taken from the radial artery every second to measure blood radioactivity using a Twilite blood sampler (Swisstrace, Zurich, Switzerland).

The functional MRI data were acquired during a 10 min time interval using a single-shot echo planar imaging sequence (300 volumes; 35 slices; repetition time, TR = 2000 ms; echo time, TE = 30 ms; flip angle, FA = 90°; field of view, FOV = 192 × 192 mm2 ; matrix size = 64 × 64; voxel size = 3 × 3 × 3.6 mm3). Diffusion-weighted images were acquired using a single-shot echo planar imaging sequence (60 slices; 30 non-colinear gradient directions; b-value = 800 s/mm² and one b=0 s/mm² image; TR = 10800 ms, TE = 82 ms; FA = 90°; FOV = 260 x 264 mm²; matrix size = 130 x 132; voxel size = 2 x 2 x 2 mm3). Additional information regarding the acquisition protocol can be found under [14].

To process the functional data, each subject’s resting state fMRI recording was pre-processed using the Configurable Pipeline for the Analysis of Connectomes (CPAC v1.4.0) [16]. This included slice-timing correction, motion correction, spatial normalization, quadratic and linear detrending of scanner drift, anatomical CompCor regression of white matter and cerebrospinal fluid activity [6], and subsequent bandpass-filtering (0.01 - 0.1 Hz). More details can be found in [14].

To process the diffusion data, structural connectomes were generated using the MRtrix3_connectome BIDS App [31], which operates principally using tools provided in the MRtrix3 package [84]. This included: DWI denoising [89], Gibbs ringing removal [40], preprocessing [1, 2], bias field correction [85], inter-modal registration [7], brain extraction [75], T1 tissue segmentation [63, 73, 75, 100], spherical deconvolution [38, 83] and probabilistic tractography [82] utilizing Anatomically-Constrained Tractography [73] and dynamic seeding [74]. The resulting fiber track files were subsequently converted into streamline counts by counting the number of streamlines that passed through one of the 400 cortical regions according to the Schaefer functional atlas[69]. In order to compensate for the bias toward longer fibers inherent in the tractography procedure, as well as differences in region size, we normalized the streamline count by the average length of the streamlines and average surface area of the two regions [34]

To process the PET data, the first 45 minutes of the PET acquisition were reconstructed offline using the NiftyPET library [52] based on the ordered subsets expectation maximization (OSEM) algorithm with 14 subsets, 4 iterations, and divided into 33 dynamic frames: 10 x 12 s, 8 x 30 s, 8 x 60 s, 2 x 180 s and 5 x 300 s. The attenuation-correction was based on the T1-derived pseudo-CT images [11]. All reconstructed PET images were motion corrected and spatially smoothed (Gaussian filter, FWHM = 6 mm). The net uptake rate constant (K*_i_*) was calculated using the Patlak plot model [64] based on the last 5 frames of the preprocessed PET images (frames between 20 to 45 minutes) and the arterial input function derived from the preprocessed arterial blood samples. The cerebral metabolic rate of glucose (*CMR_glc_*) was calculated by multiplying the K*_i_* map with the concentration of glucose in plasma of every participant, divided by a lumped constant of 0.65 [93]. Finally, the *CMR_glc_* maps were partial volume corrected using the GM, WM and CSF masks derived from the T1 images using the iterative Yang method [94] and finally registered to the MNI152NLin6ASym 3mm template through the anatomical image.

#### Additional data

In addition to the metabolic map of glucose uptake from the TUM dataset, we complemented our analysis with a normative, group-average map of oxygen metabolism (*CMR_O_*_2_) from [86], acquired through neuromaps [50]. The brain map was derived from 33 healthy, right-handed neurologically normal participants (mean age 25.4 ± 2.6 years, 58% females) that were recruited from the Washington University community. The recording was performed using a Siemens model 961 ECAT EXACT HR 47 PET scanner (Siemens/CTI) with 47 slices encompassing an axial field of view of 15 cm. Transverse resolution was 3.8–5.0 mm FWHM and axial resolution was 4.7–5.7 mm full width at half maximum (FWHM). Attenuation data were obtained using 6868GeGa rotating rod sources to enable quantitative reconstruction of subsequent emission scans. Emission data were obtained in the 2D mode (interslice septa extended). The PET data were reconstructed using a ramp filter (*≈*6 mm FWHM) and then blurred to 12 mm FWHM. Distribution of *CMR_O_*_2_ was measured with a 40 second emission scan (derived from a 120 second dynamic scan) after brief inhalation of 60 mCi of [15O]oxygen in room air. More details on the recording parameters and processing can be found in [86].

### Network control theory

Network control theory (NCT) is a framework that provides the approach to computing metrics related to the control of brain activity.

Fundamental to NCT is the definition of the brain as a networked system with nodes connected by edges and defined by an adjacency matrix *A*. Biologically, the matrix *A* represents the thick bundles of myelinated axonal fibers that run through large-scale connections of brain regions [77]. These fibers are thought to play a critical role in coupling the activity of distant brain regions [3]. By accounting for connections of at the brain-region level, an equation relating brain structure to time-evolving brain activity **x**(*t*) can be formulated as:

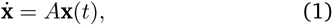

where **ẋ** and **x** are of size *N* x1 and *A N* x*N* , with *N* is the number of brain regions. In practice, the matrix *A* is normalized in order to avoid infinite growth, and as such defined as:

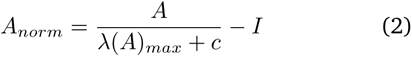

where *λ*(*A*)*_max_* is the largest eigenvalue of *A* and *c* a constant that determines if the system approaches the corresponding mode of *A* (*c* = 0) or decays towards zero (*c >* 0). Here, c is set to 1. For simplicity, *A_norm_*will be referred to as *A* from now on, assuming that all the following equations make use of a normalized matrix.

Looking at the assumptions made, equation 1 imposes that the temporal evolution of the brain is a *linear* function that is described by its connectivity and its current state in time. In addition to linearity it is important to note that this equation assumes that *A* is not changing, i.e., the brain is a time-invariant system, and that brain dynamics are noise-free [39, 78].

From here, equation 1 can be extended to account for controlled dynamics that steer the system away from its natural trajectories through an external input. Formally,

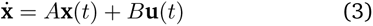

with **u**(*t*) representing the injected control input into the system at time *t* and the columns in *B* representing the total number of voxels or regions of interest to control. Depending on the number of regions *m* that the modeler decides to control, **u**(*t*) is of size *m*x1 and, correspondingly, *B* is of size *N* x*m*. Each element in *b_ij_* represents the influence that an external input *u_j_*(*t*) has on region *i*, where *j* = 1*,.., m* and *i* = 1*, …, N*. By including controlled dynamics in equation 3 it is now possible to model optimal trajectories between brain states and calculate their optimal control energy.

#### Optimal control energy

Optimal control energy provides a measure of the controllability of a system in terms of 1) the state trajectory traversed and 2) the work required to reach a state. It further specifies a time horizon *T* during which the control input *u*(*t*) is effective in moving the system from the initial state *x*(0) = *x*_0_ to the goal state *x*(*T*) = *x_T_*.

Formally, the problem of reaching *x_T_* starting from *x*_0_ while keeping 1) and 2) minimal is as follows:

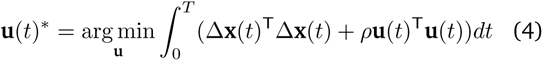

with Δ**x**(t) being the distance to the goal state at time *t*, i.e.,

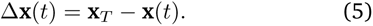

Moreover, *ρ* determines the relative weighting between the costs associated with the length of the state trajectory and input energy. An usual choice is to set *ρ* = 1 as to weight both goals equally [32, 39]. Once equation 4 is minimized, an optimal control energy can be derived from **u**(*t*)*^∗^*:

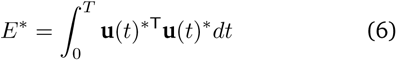

The optimal energy *E^∗^* was computed for each spatial unit (in this case regions from a parcellation). These values therefore represent the quadratic input across model time to optimally move towards a goal state given an initial state.

Readers interested in a more in-depth discussion about the mechanisms of network control theory are invited to read the review from [78], as well as [60]. For more discussions on parameter sensibilities of the model, interested readers are referred to the exhaustive study by [39].

### Spatial autocorrelation preserving null model

For all of our hypothesis tests that relied on comparing two brain maps, we employed BrainSMASH [12] as implemented in the neuromaps [50] library. Briefly, the methodology of BrainSMASH entails two steps: 1) random permutation of values within a designated brain map and 2) smoothing and re-scaling to restore the spatial autocorrelation characteristic of the original data.

In the first step, values within the brain map undergo random permutation. Subsequently, the permuted data undergoes a transformation to reintroduce spatial autocorrelation. This transformation is expressed as

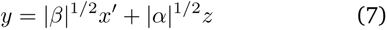

where *x^′^* represents the permuted data, *z ∼ N* (0, 1), and *α* and *β* are estimated via least-squares optimization, aligning variograms of the original and permuted data.

### Repeated measures correlation

Repeated measures correlation (*r_rm_*) is a statistical technique developed by Bakdash et al. [4] to examine the common association between variables within each subject, focusing on accounting for within-individual associations in paired measures. This method is particularly valuable when dealing with repeated measures on the same participants — in our case four control cost estimates for each hierarchical transition. Because this method only assesses the common slope across participants, we can investigate whether there is a consistent relationship trend throughout measurements while automatically correcting for individual confounding factors such as the participant’s gender or age.

## Acknowledgments

We thank Filip Milisav, Justine Hansen, Vincent Bazinet, Asa Farahani, Zhen-Qi Liu, Moohebat Pourmajidian, Laura Suarez, and Markus Ploner for helpful discussion. EGC acknowledges support from the Molson Foundation, the IFI fellowship of the German Academic Exchange Service and the Elite Network of Bavaria. AIL was supported by the Natural Sciences and Engineering Research Council of Canada (NSERC), [funding reference number 202209BPF-489453-401636, Banting Postdoctoral Fellowship]. VR has been funded by the European Research Council (ERC) under the European Union’s Horizon 2020 research and innovation program (ERC Starting Grant, ID 759659).

---

**Figure S1.**
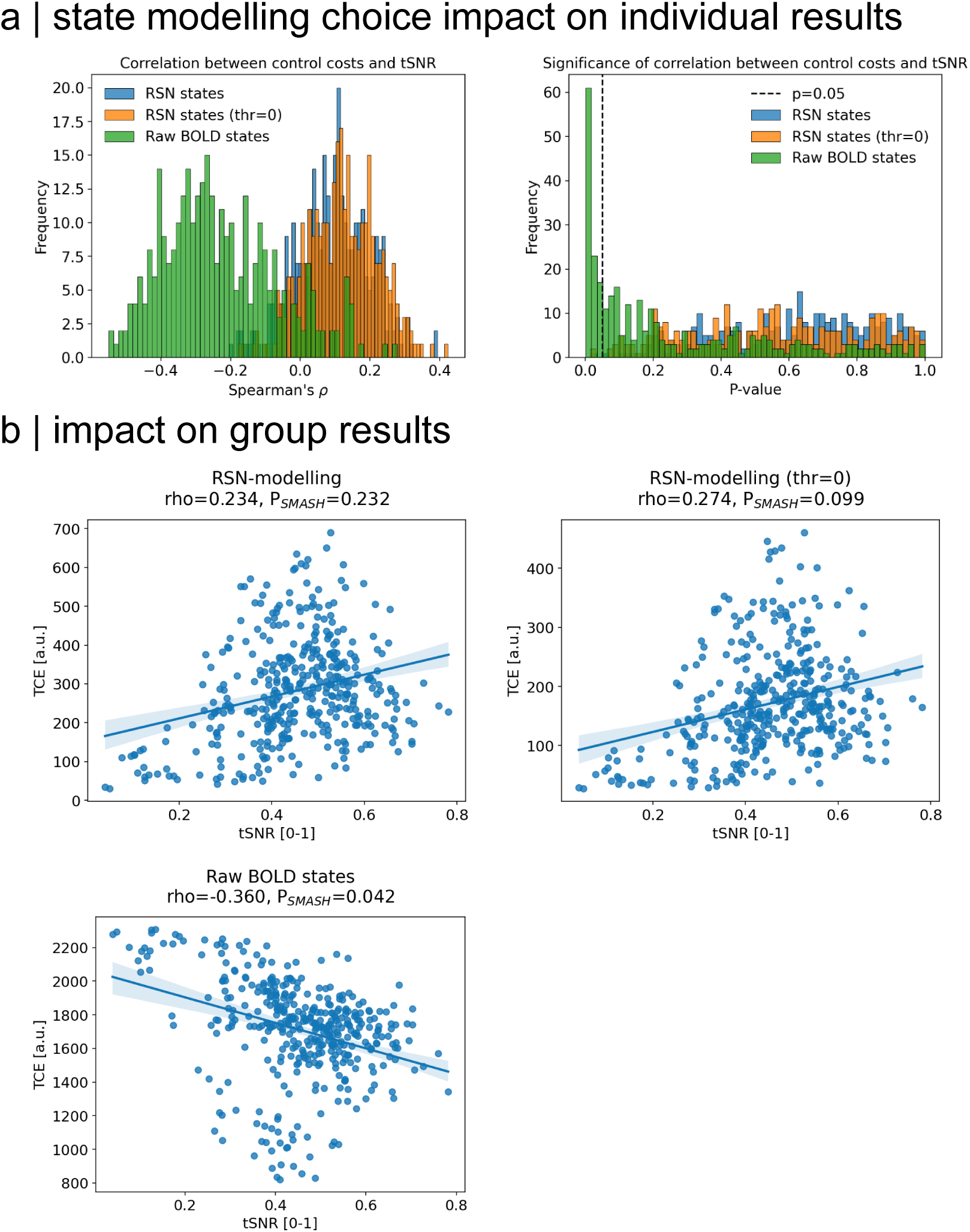
Intrinsic network modelling validation. (a) The temporal signal-to-noise ratio (tSNR) of each individual’s recording was estimated by dividing their mean signal prior to demeaning with their signal variability [92]. This metric was compared to their control costs from three modelling approaches: 1) our reported approach to use intrinsic networks as states, where only time points with an average network activity above 0.5 are selected; 2) same as 1) but with an activity threshold of 0; 3) raw BOLD activity are used as states. The first two approaches result in TCE maps that correlate similarly to their respective tSNR (left), with only few subjects showing significance in their associations(right). Using the third approach, however, results in control costs being negatively correlated to tSNR (left), and more individual’s control maps being significantly related to the tSNR of their recording (right). (b) We see similar results on a group-level, where we average all control maps and tSNR across subjects. In both state modelling approaches, we observe that control costs and tSNR are not significantly associated, however, the lower threshold variant results are statistically more unlikely to be discernible [Spearman *ρ* = 0.234, *PSMASH* = 0.232 for thr=0.5 and Spearman *ρ* = 0.274, *PSMASH* = 0.099 for thr=0]. Critically, the group-average map of TCE significantly correlates with tSNR when opting for no state modelling [Spearman *ρ* = -0.36, *PSMASH* = 0.042].

**Figure S2.**
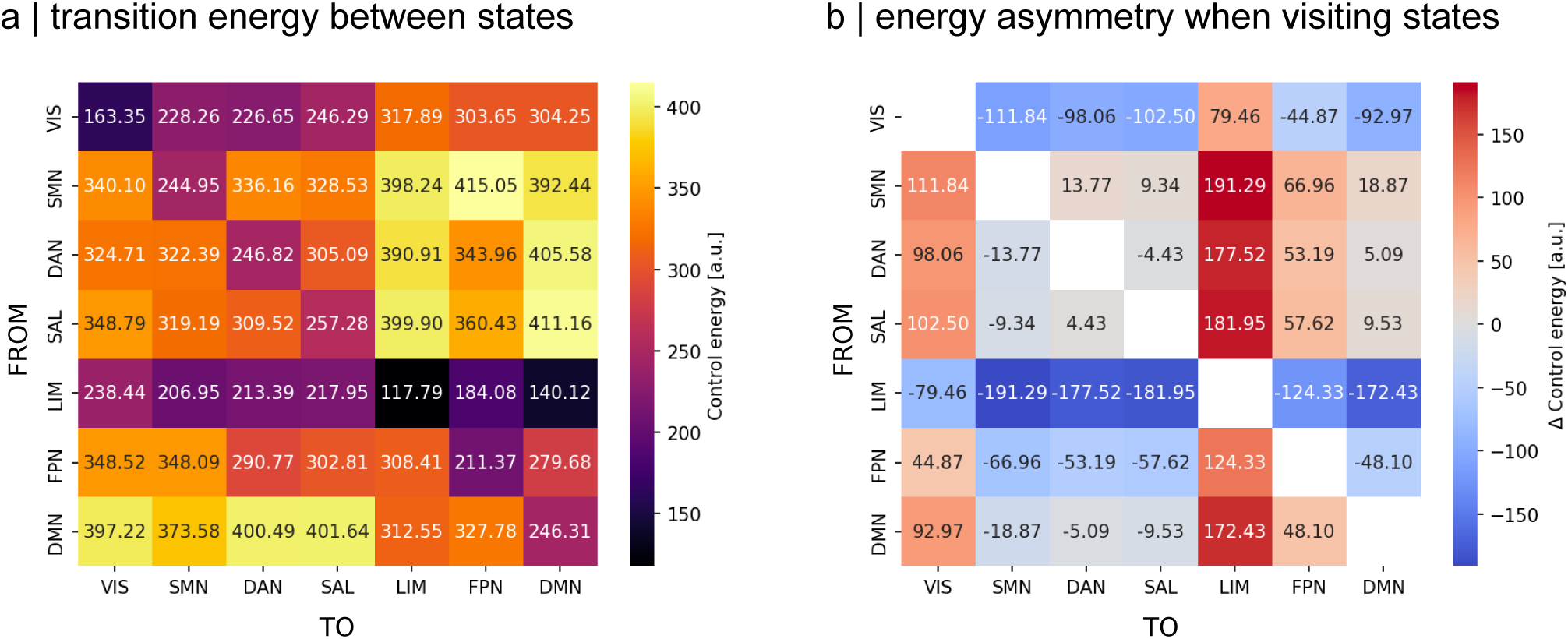
Transition energy matrix. (a) We simulate the control energy to transition between all of pairs of intrinsic networks. Similar to previous work [48, 61], we find that the costs to transition into a state are dissimilar to the costs of leaving it. (b) We highlight this asymmetry by subtracting the transition energy matrix with its transpose, thereby accentuating the difference between going into and leaving a state.

**Figure S3.**
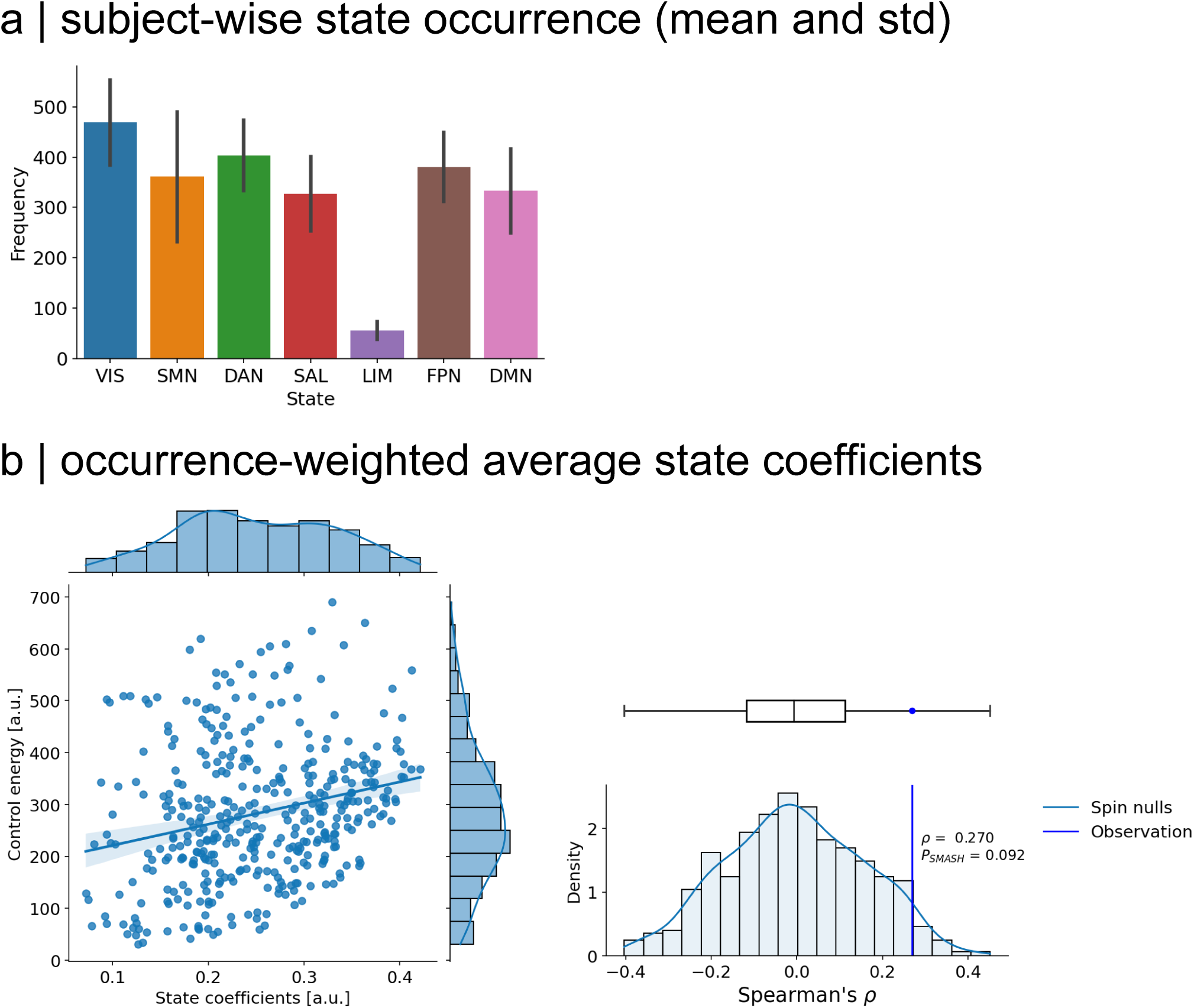
Control costs vs. state coefficients. (a) State occurrences are counted for each individual. We report the average and standard deviation of each state distribution. (b) Left: Weighting each network map by their occurrence and then summing all maps yields an occurrence-weighted state coefficient map. This is averaged across all individuals and compared to the group-level TCE map. Right: TCE and state coefficients are not significantly correlated when accounting for spatial autocorrelation [Spearman *ρ* = 0.27, *PSMASH* = 0.092].

**Figure S4.**
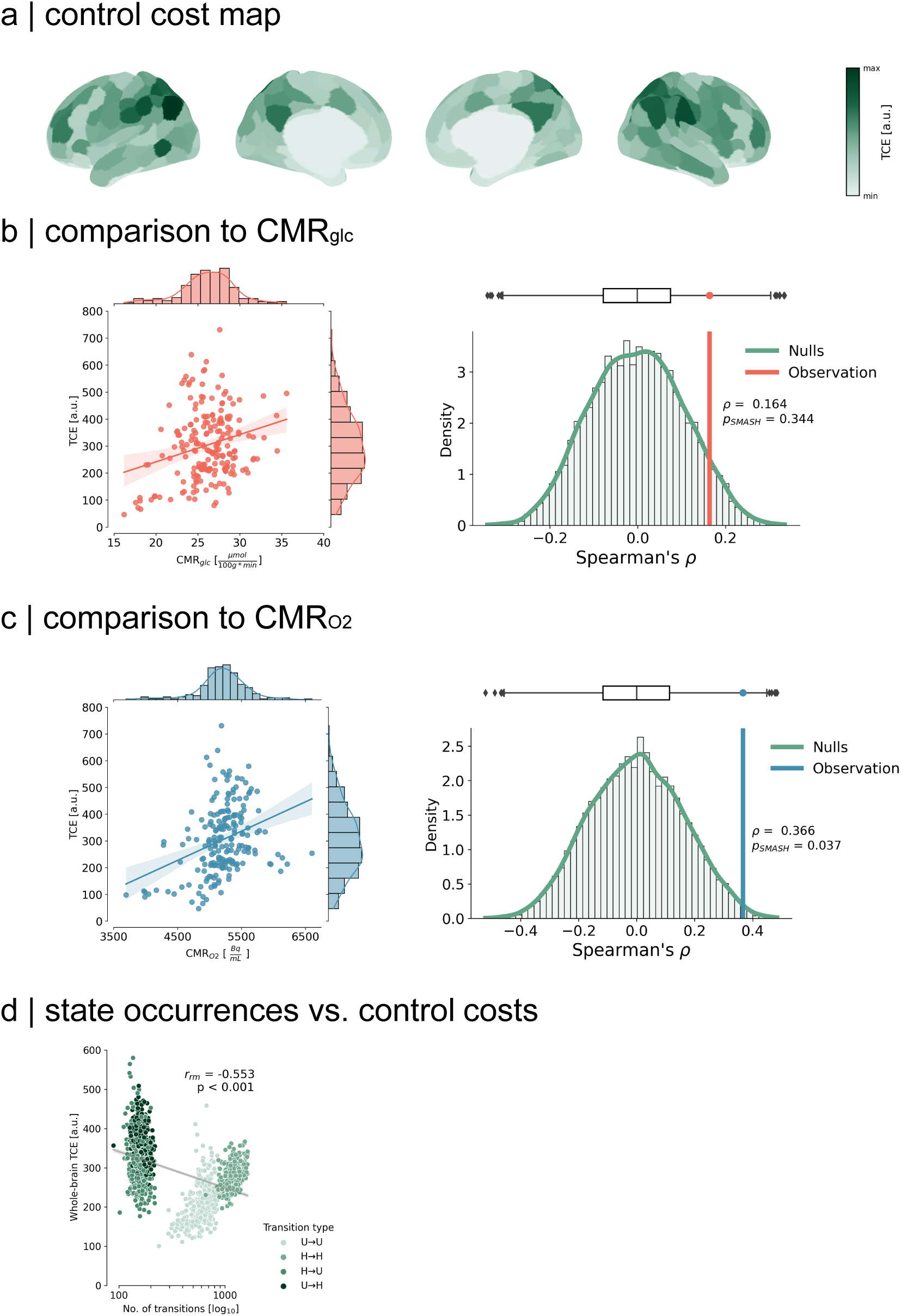
Replication with Schaefer 200 parcellation|. (a) TCE map across both hemispheres. (b) No significant relationship to *CMR_glc_* [Spearman *ρ* = 0.164, *P_SMASH_* = 0.344]. (c) Significant relationship to *CMR_O_*_2_ [Spearman *ρ* = 0.366, *P_SMASH_* = 0.037]. (d) Whole-brain TCE is inversely related to the number of transitions across hierarchies [*rrm* = -0.553, P < 0.001].

**Figure S5.**
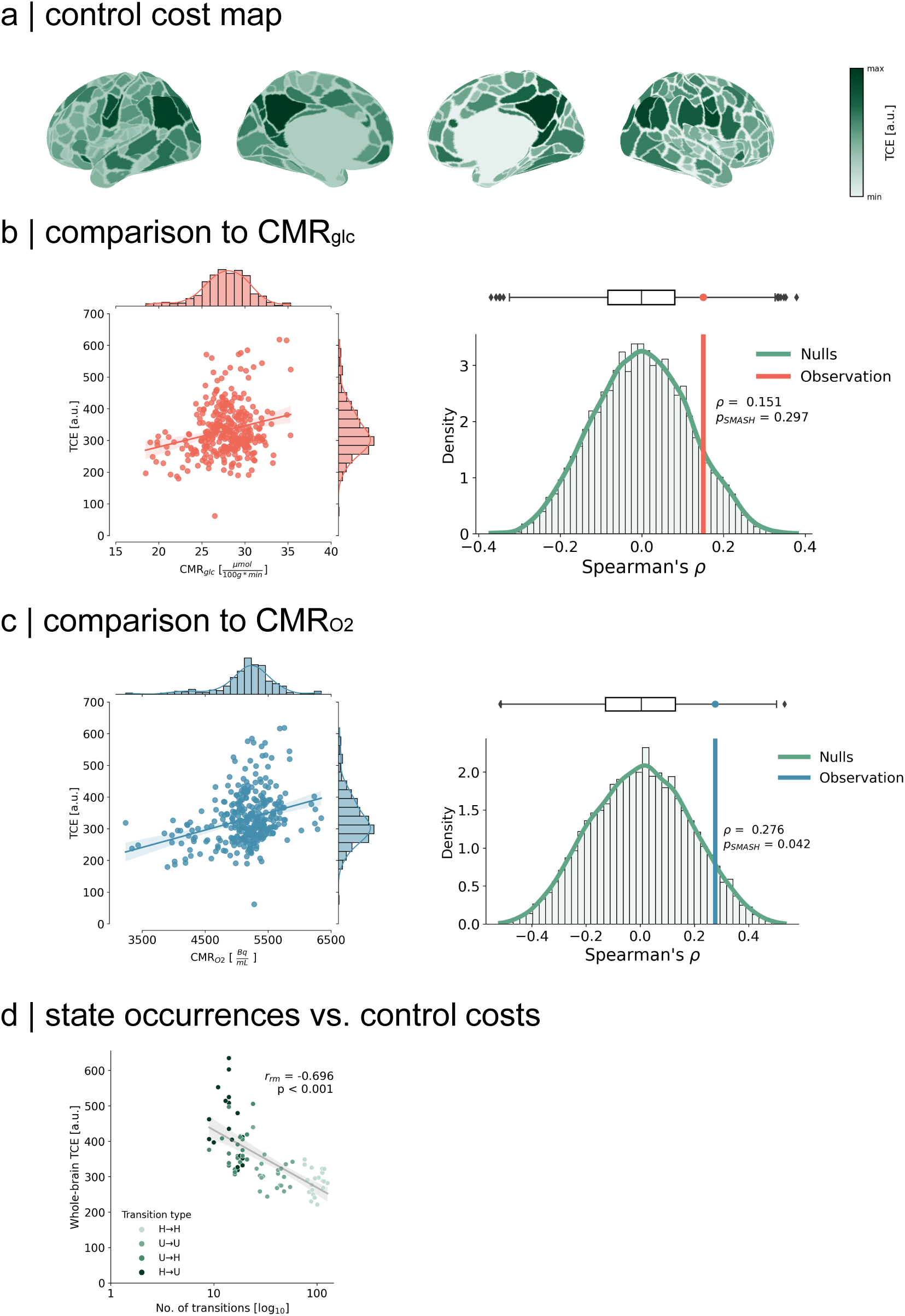
Replication with Gordon parcellation. (a) TCE map across both hemispheres. (b) No significant relationship to *CMR_glc_* [Spearman *ρ* = 0.151, *P_SMASH_* = 0.297]. (c) Significant relationship to *CMR_O_*_2_ [Spearman *ρ* = 0.276, *P_SMASH_* = 0.042]. (d) Whole-brain TCE is inversely related to the number of transitions across hierarchies [*rrm* = -0.696, P < 0.001].

**Figure S6.**
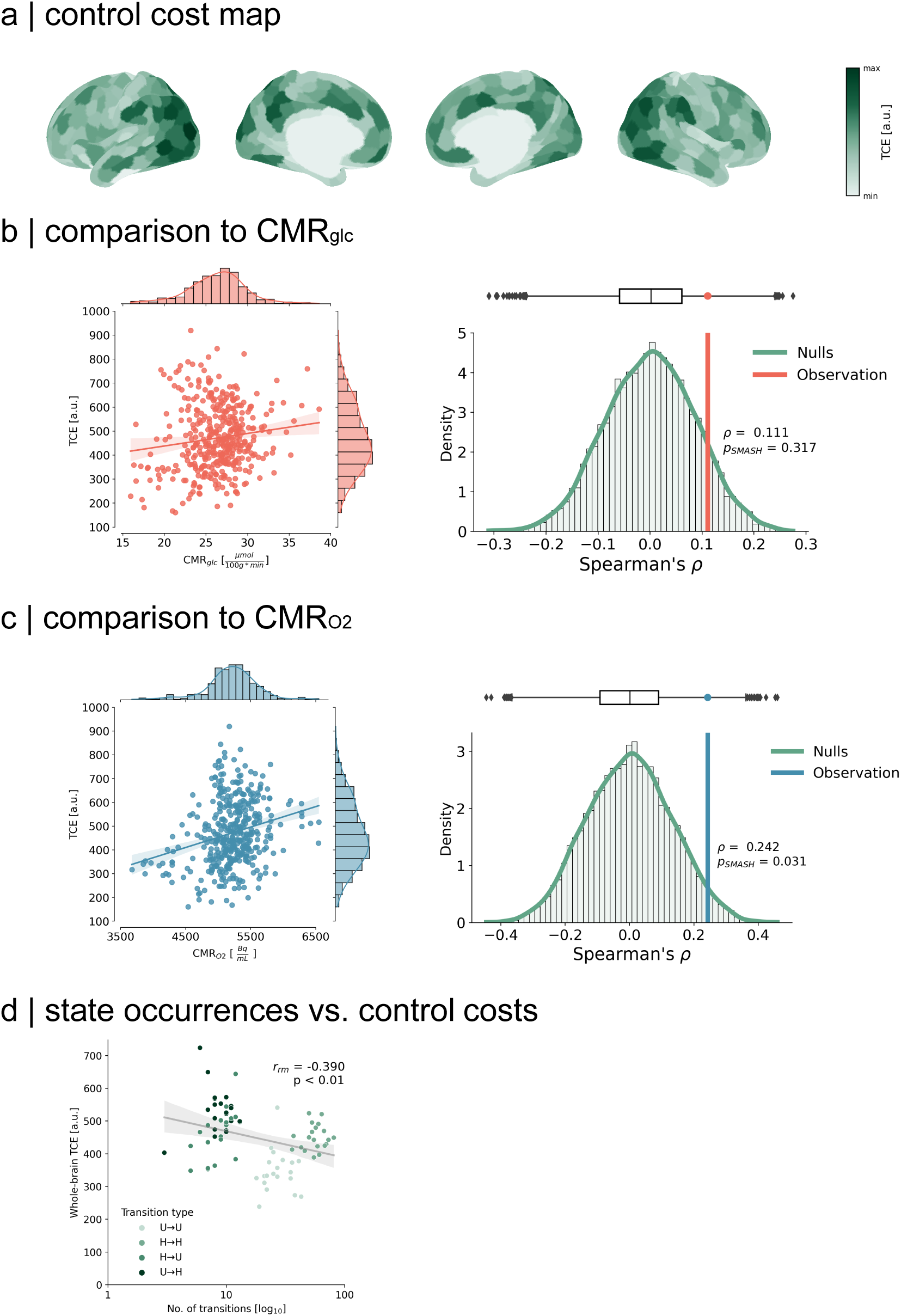
Replication with separate dataset. [14] **|** (a) TCE map across both hemispheres. (b) No significant relationship to *CMR_glc_* [Spearman *ρ* = 0.111, *P_SMASH_* = 0.317]. (c) Significant relationship to *CMR_O_*_2_ [Spearman *ρ* = 0.242, *P_SMASH_* = 0.031]. (d) Whole-brain TCE is inversely related to the number of transitions across hierarchies [*rrm* = -0.39, P < 0.01].

